# Limited emergence of resistance to Integrase strand transfer inhibitors (INSTIs) in HIV-experienced patients failing dolutegravir-based antiretroviral therapy: Cross-sectional analysis from a Northeast Nigerian cohort

**DOI:** 10.1101/2022.11.08.515598

**Authors:** Adam Abdullahi, Ibrahim Musa Kida, Umar Abdullahi Maina, Amina Husaini Ibrahim, James Mshelia, Haruna Wisso, Abdullahi Adamu, James Ezenwa Onyemata, Haruna Yusuph, Sani H. Aliyu, Man Charurat, Alash’le Abimiku, Lucie Abeler-Dorner, Christophe Fraser, David Bonsall, the PANGEA consortium, Steven A. Kemp, Ravindra K. Gupta

## Abstract

**Background:** Owing to high levels of resistance to previous first-line non-nucleoside reverse transcriptase inhibitors (NNRTI)-based antiretroviral therapy (ART), consolidated recommendations since 2019, from the WHO and others, have indicated that dolutegravir (DTG) is the preferred drug of choice for HIV treatment, globally. There is a paucity of resistance outcome data from non-B HIV subtypes circulating across West Africa.

**Aim:** We aimed to characterise the mutational profiles of HIV-positive patients from a small North-East Nigeria cohort, failing a DTG-based ART regimen.

**Methods:** Plasma samples were collected and stored from 61 HIV-1 infected participants. Following failure of DTG-based ART, all samples were sequenced by Illumina whole-genome, ultra-deep sequencing. Sequencing was successful in (n=33) participants with median age of 40 years and median time on ART of 9 years. HIV-1 subtyping was performed using SNAPPy. Haplotype reconstruction and transmission were inferred using standard phylogenetic methods.

**Result:** Most patients had mutational profiles that were reflective of prior exposure to first- and second-line ART including exposure to thymidine analogues, efavirenz and nevirapine. One patient had evidence of major INSTI DRMs (T66A, G118R, E138K and R263K), reducing efficacy of DTG. The participant was aged 18, infected with a subtype G virus and likely vertically infected.

**Conclusion:** This study found low level resistance to DTG in the cohort, with one patient having high-level resistance to DTG and other INSTIs. Critical population level and long-term data on DTG outcomes are required to guide implementation and policy action across the region.

## Introduction

In the context of rising pre-treatment NNRTI drug resistance^1,2^, the World Health Organisation (WHO) recommended dolutegravir (DTG) as the preferred antiretroviral therapy (ART) drug of choice for both newly diagnosed and individuals transitioning from previous regimens^3^. Safety, potency, tolerability and cost-effective characteristics of dolutegravir (DTG) supported this change^4^ subsequently, countries across sub-Saharan Africa (SSA) have consequently rolled-out dolutegravir as part of standard treatment. Roll out across the region started in 2019 and is expected to continue, aided by the availability of a low-cost, generic fixed-dose co-formulation of tenofovir, lamivudine and dolutegravir (TLD)^5^.

Dolutegravir-based antiretroviral therapy (ART) have been commercialised and sold in Nigeria since 2019 with the national treatment guideline recommending transitioning to DTG-based ART in both virally suppressed and unsuppressed patients since 2020^6^. There is no indication of virological or resistance testing prior to transitioning to DTG-based ART and therefore, majority of patients transitioned without prior viral load or resistance testing. Data from the ADVANCE and NAMSAL clinical trials^7,8^, which recruited ART naïve participants exclusively in SSA showed no evidence of emergence of drug resistance mutations (DRMS) on DTG-based ART. Data from treatment experienced patients transitioning to TLD is limited although data is starting to emerge.

Given the high proportion of treatment experienced HIV patients with resistance following failure of previous first and second-line ART^9^, data on resistance outcome following failure of DTG in non-B subtypes is highly valuable. Here, we present data on drug resistance using Next-Generation Sequencing (NGS) in a small Nigerian cohort failing DTG-based ART following roll-out.

## Methods

### Study population and design

This study was a cross-sectional study performed at the University of Maiduguri Teaching Hospital, Borno State, Nigeria between January, and June 2021. Study criteria included participants who were failing a DTG-based ART, ≥18 years of age and attending routine clinic visits. We defined virological failure as two consecutive HIV-1 RNA > 1000 copies/ml following exposure to a DTG-based ART for 6 months. Patients who met inclusion criteria voluntarily signed informed consent. Available demographic data including age, gender, ART regimen, duration on ART and current CD4 count were collected from clinical files and recorded in Microsoft excel.

### Laboratory methods

Plasma was separated from whole venous blood in EDTA within 2 hours of collection and stored immediately at −80°C. Plasma viral load testing and CD4 count were performed at the Defence Reference Laboratory, Asokoro Abuja using the COBAS AmpliPrep/COBAS TaqMan HIV type 1 (HIV1) v2.0 test (Roche Diagnostics, Basel, Switzerland). Whole genome sequencing of 61 blood samples was performed according to the Bonsall et al protocol^10^. Briefly, total RNA was extracted from HIV-infected plasma samples, washed in ethanol, and eluted using the NUCLISENS easyMAG system (bioMérieux). Libraries were prepared using the SMARTer Stranded Total RNA-Seq Kits v2 (Clontech, Takara Bio) according to the manufacturer protocol. Total RNA was denatured, and reverse transcribed to cDNA and a total of 500ng of pooled libraries were hybridised to custom HIV-specific biotinylated 120-mer oligonucleotides (xGen Lockdown Probes, Integrated DNA Technologies). Captured libraries were then PCR amplified to produce a final pool for sequencing with an Illumina MiSeq (San Diego, CA, USA) to produce up to 300-nucletoide paired-end reads. FastQ files were trimmed of adapters and mapped iteratively to the best available reference from a curated alignment of 3000 HIV-1 genomes with SHIVER^11^. Resistance genotyping was performed using an in-house script that determines the prevalence of DRMS in each sequencing reads and calculates an overall score (1-4) for each ART, according to the Stanford HIV drug resistance algorithm (v9.1).

### Bioinformatics analysis

Haplotypes were reconstructed using CliqueSNV v2.0.3^12^. Phylogenies were inferred with IQTREE2 v2.2.2^13^ using a GTR+F+I+R4 model with 1000 rapid bootstraps. Inference of transmission was made Phyloscanner v1.8.1^14^ using overlapping windows of 150 bp across the whole genome. Phylogenies were rooted on a HIV-1 subtype G consensus sequence downloaded from the Los Alamos National HIV Database. HIV-1 subtyping was performed using SNAPPy v1.0. Prediction of co-receptor usage was made using TROPHIX (prediction of HIV-1 tropism). Available at: http://sourceforge.net/projects/trophix/).

### Statistical analyses

The characteristics of the study population were summarized as either categorical or continuous variables and reported as either proportions or medians with interquartile ranges (IQRs), respectively. Analyses were performed with STATA v17 (StataCorp, College Station, TX).

### Ethics

The study was approved by the University of Maiduguri Teaching Hospital Ethics Committee (UMTH/REC/21/714). All participants provided written informed consent.

## Results

61 had samples available for resistance testing. Following sequencing and quality control, 33 samples (61.7%) were of sufficient quality (i.e., fully in-tact pol gene and depth of 500x) to determine DRMS and minority variants (Median viral load in these samples was 4.1 log10 copies/ml (range 3.0-5.2). Using a minimum threshold of 20%, NRTI, NNRTI, PI and INSTI DRMS occurred in 17 (50%), 24 (70.6%), 4 (11.8%) and 1 (2.9%) of samples respectively (Figure 1a). Dual-class resistance occurred in 17 (50%) patients and tri-class mutations occurred in 5 (14.7%) patients. Consistent with likely long-term exposure to lamivudine, the most prevalent NRTI mutation DRMS was M184V. The most prevalent NNRTI DRMS was K103N, reflecting previous exposure to nevirapine and efavirenz. Mutational profiles were similar across 2, 10 and 20% interpretative thresholds (Figure 1b).

**Figure 1a:**
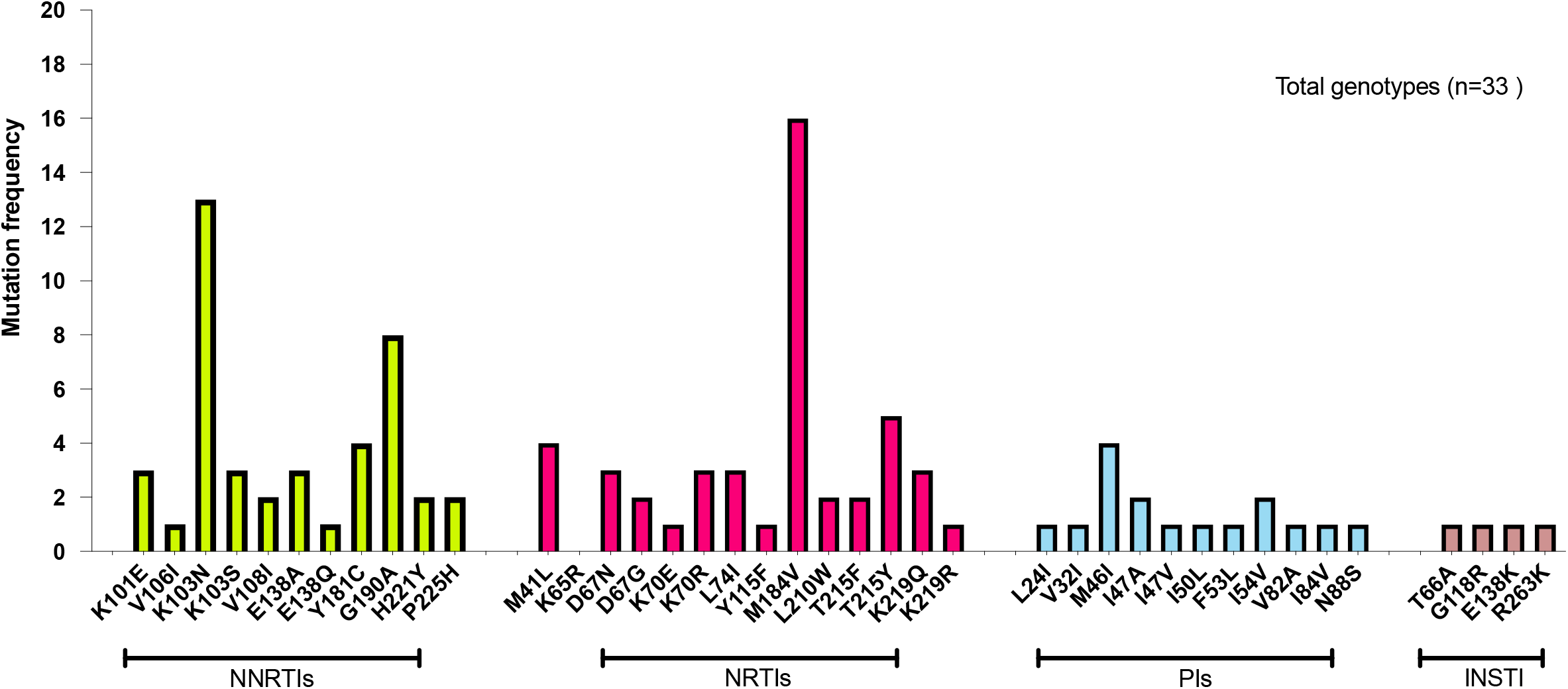
Proportion of patients with resistance associated mutations using the Stanford algorithm.

**Figure 1b:**
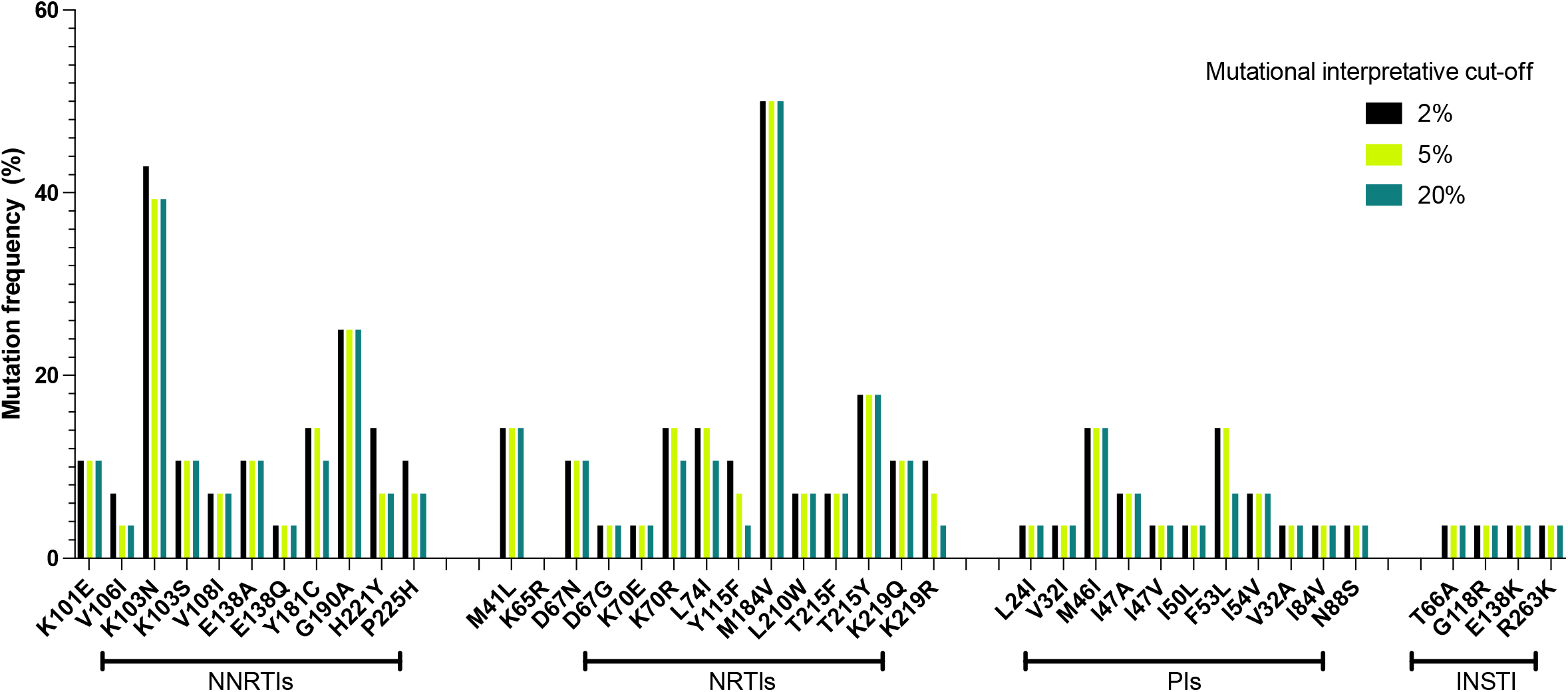
Proportion of patients with resistance associated mutations using the Stanford algorithm, sub-divided into interpretational cut-offs of 2, 5 and 20%. Evidence suggest that minority variants may play a role in drug resistance (https://doi.org/10.1128/mbio.00269-22).

One patient of interest was found to have high-level resistance to NRTIs, NNRTIs and INSTIs, including almost complete resistance to the novel long-acting injectable, cabotegravir. Mutations included inE138K, inG118R, inT66A, inR263K, rtH221Y, rtV108I, rtK103N, rtM184V, rtM41L, rtA98G and rtT215Y, all at frequencies of >40%, with a mean read depth of 770x. This patient was established on DTG for a median of 1.2 years and on ART for a median of 12 years. Clinical data on other clinical data including nadir CD4 counts were unavailable. Using the SNAPPy HIV-1 subtyping tool, almost 40% of viruses were assigned to Subtype G, and 15% were G_A1 subtypes. The remaining viruses were recombinants (Table 1).

**Table 1:**
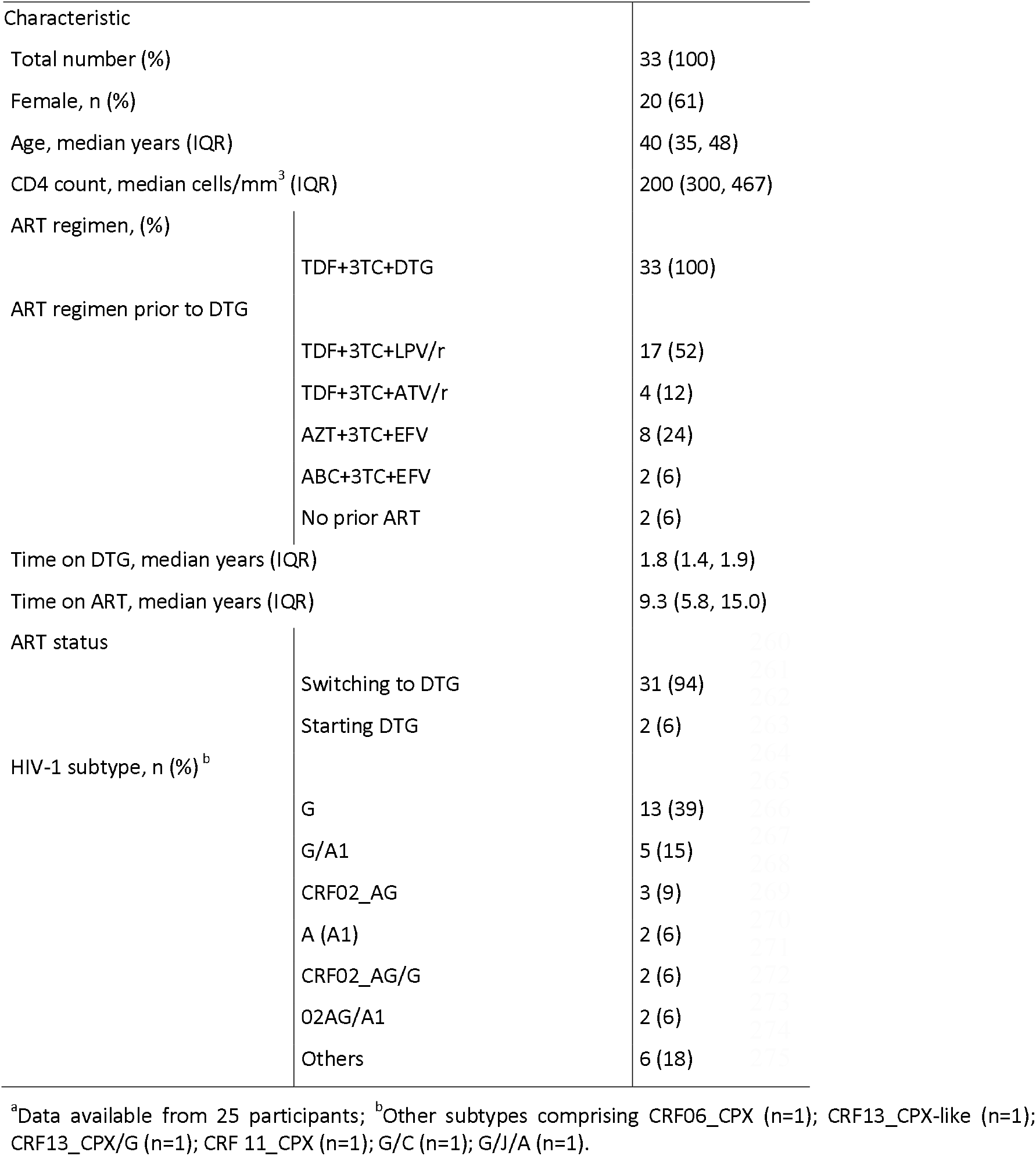
Characteristics of study participants with successful genotyping

To identify within-host diversity and potential transmission of DRMs between patients, we reconstructed viral haplotypes (Figure 2) for each patient. These were homogeneous and the same resistance mutations were identified on all reconstructed haplotypes for each patient. Following this, we investigated whether there was evidence of direct transmission between any patients in this cohort (Supplementary Figure 1). However, no significant transmission pairs were identified, indicating that there were several intermediaries between patients which have not yet been sampled. Of note, two patients’ virus was predicted to use the CXCR-4 co-receptor.

**Figure 2.**
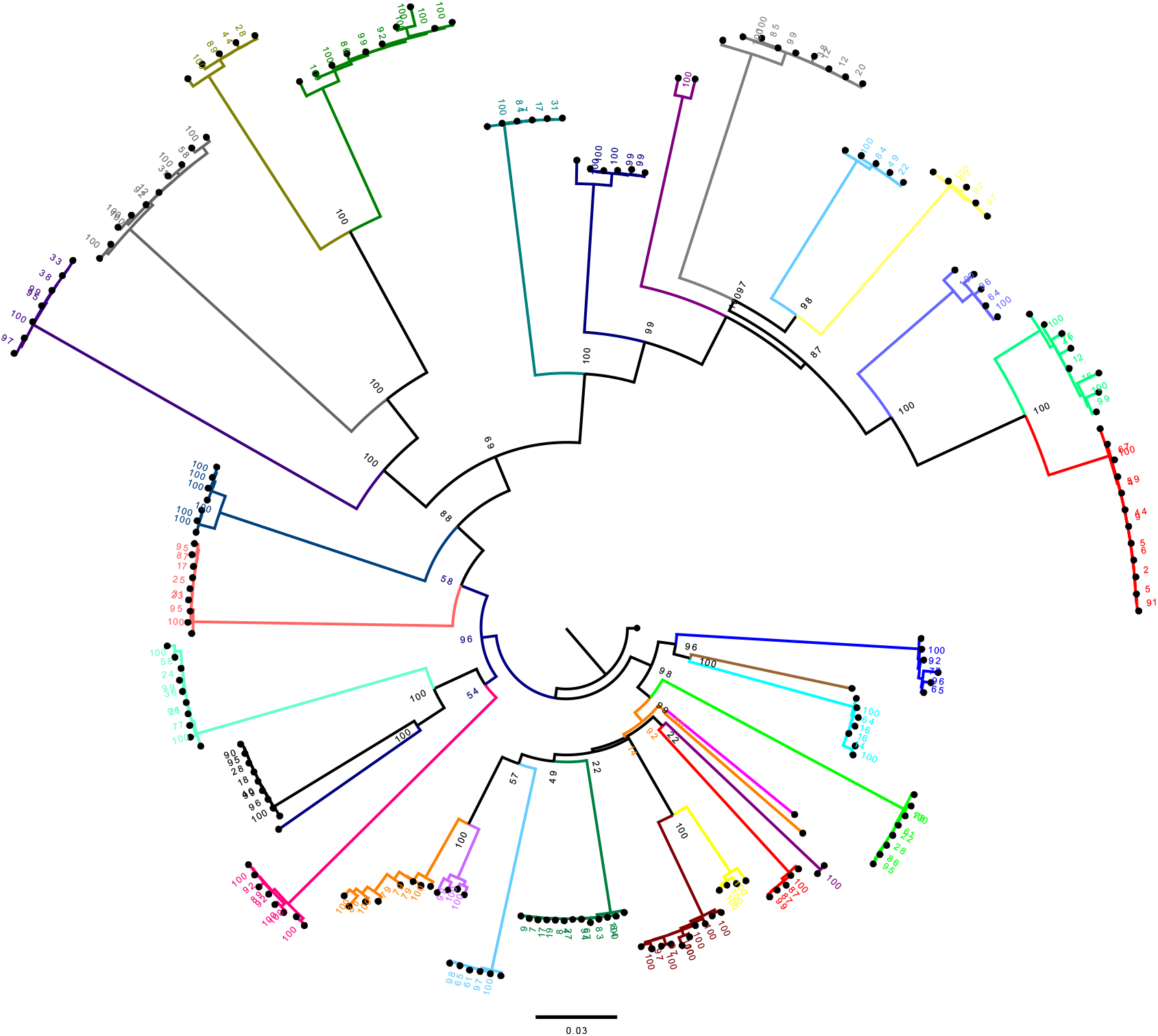
Maximum-likelihood phylogeny of reconstructed haplotypes from each patient. Haplotypes were homogeneous and had the majority of the same mutations on each haplotype. Bootstraps are indicated at each node.

## Discussion

After rollout of previous first-line NNRTI based ART, DRMS were previously observed within first year of failure in around 15 and 35% of patients with resistance emerging against both lamivudine, tenofovir and NNRTIs^16,17^. Drug resistance has been associated with mortality in hospitalised individuals in LMIC settings^18^. The second generation INSTI DTG has been systematically adopted and rolled out across the SSA region since 2019, with concerns of the emergence of resistance associated mutations following failure under a limited monitoring infrastructure. In a recent analysis of pooled evidence on virological and resistance outcomes following DTG failure in SSA region, there was an overall high rate of virological response to DTG; 88.5% (95% CI: 73.8-97.8) with the overall proportion of patients failing showing limited evidence of DRMS^19^ over short periods of time. It is likely that prolonged virologic failure will select for DRM to components of ART regimens within the viral quasispecies as a result of intrahost evolution^20,21^.

It is important to note that pre-existing DRMS prior to switch to DTG may be critical to both virological and resistance outcomes with study evidence suggesting pre-existing NNRTI mutations reducing the short term efficacy of DTG^22^, although other studies across the region have shown similar rates of both virological and resistance outcomes in ART naïve^23,24^ and experienced patients^25,26^ (with no evidence of DRMs) and ART experienced patients with historical evidence of NRTI mutation (especially M184V/I)^27–29^.

Here, in this cross-sectional analysis, we assessed drug resistance using next-generation sequencing in a small cohort of HIV-1 infected subjects failing DTG-based ART using a failure threshold of 1000 copies/ml. Most patients were treatment experienced and amongst 33 participants with sequence data, mutational patterns observed were reflective of exposure to previous first-line NNRTI with only 1/34 (3%) showing evidence of DRMS against DTG or protease inhibitors. The individual with DTG resistance was vertically infected with evidence of selection of mutations conferring high level resistance to dolutegravir and other INSTIs i.e T66A, G118R, E138K (accessory) and R263K. The T66A mutation is non-polymorphic and primarily selected by elvitegravir (EVG) and raltegravir (RAL) with ∼9-fold reduction in susceptibility to EVG but minimal impact on other INSTIs whilst the E138K mutation has negligible effect on susceptibility to INSTIs although a combination of E138K and other DRMs may lead to further decreased susceptibility to DTG^30^. Further, the G118R and R263K mutations observed in this patient, which causes between 2 to 15 fold reduction to DTG susceptibility^31,32^ have also been observed in patients experiencing virological failure to INSTI drug agents in non-B HIV subtypes^33–36^. It is likely that the G118R mutation emerged first and led to the accumulation of other mutations including the E138K compensatory mutation as the G118R mutation has been described as the DTG-resistance pathway in non-B subtype^31,37^.

In context of the continuously expanding use of DTG, the most convenient approach to managing patients on DTG with persistent viraemia remains uncertain especially in resource limited setting such as this, where drug resistance testing capacity remains limited^38^. Several factors may increase the likelihood of the emergence of DTG resistance across the region including prolonged virological failure due to lack of routine virological monitoring^39,40^and poor treatment adherence which is an independent determinant of virological outcome^41^ in these settings. Further analyses of resistance across SSA are warranted over extended periods, as well as surveillance for INSTI resistance in newly diagnosed individuals. This is even more critical given INSTI based long acting injectables are being considered as PreP^42^.

## Acknowledgements

We are thankful to the volunteers who participated in the study. A.A. is supported by Africa Research Excellence Fund Research Development Fellowship (AREF-318-ABDUL-F-C0882) and Cambridge-Africa award. A.A. and S.A.K. are supported by Bill and Melinda Gates Foundation via the Phylogenetics and Networks for Generalised Epidemics in Africa (PANGEA) (grant number OPP1175094). R.K.G. is supported by a Wellcome Trust Senior Fellowship in Clinical Science (WT108082AIA).

## Author contributions

Study conception, design and administration: A.A, R.K.G and I.M.K; data collection: A.A, I.M.K, A.H.I, J.M, A.Ad, J.E.O, H.Y, S.H.A, A.Ab, S.K and R.K.G; data analysis: A.A., I.M.K, S.K, A.Ab and R.K.G; data interpretation: A.A, S.K, A.Ab and R.K.G; manuscript preparation; A.A wrote the first draft of the manuscript with the critical input of all co-authors. All authors reviewed the results and approved the final version of the manuscript.

**Supplementary Figure 1:**
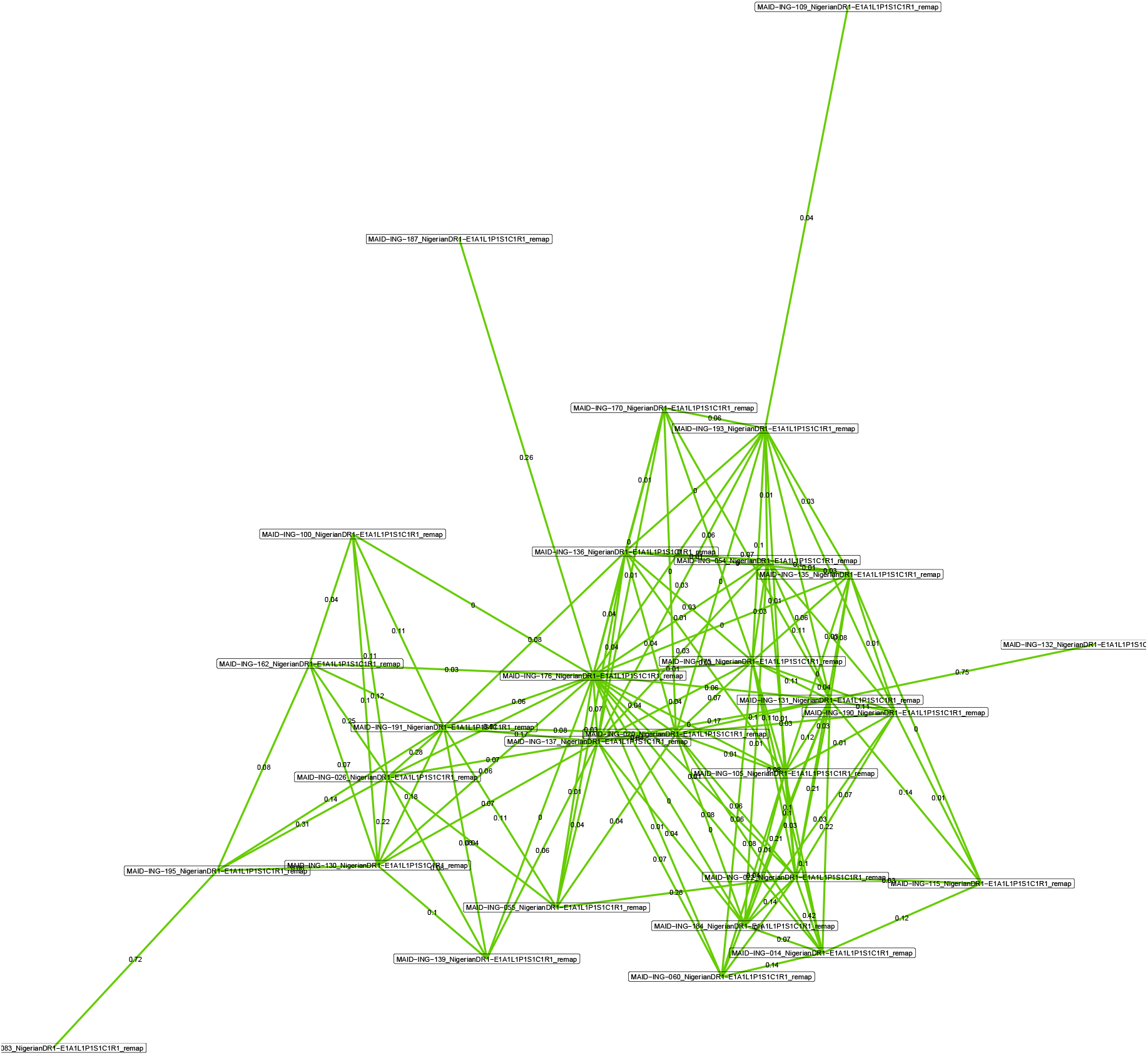
Inferred transmission network of all patients in the cohort. Green lines indicate a degree of linkage between two sequences, but without sufficient statistical support to indicate a direct transmission event has occurred.

